# Laser Ablation of the Pia Mater for Insertion of High-Density Microelectrode Arrays in a Translational Sheep Model

**DOI:** 10.1101/2020.08.27.269233

**Authors:** Kevin M. Boergens, Aleksandar Tadić, Matthew S. Hopper, Ingrid McNamara, Kunal Sahasrabuddhe, Yifan Kong, Malgorzata Straka, Harbaljit S. Sohal, Matthew R. Angle

**Affiliations:** Paradromics, Inc., Austin, TX, USA

## Abstract

The safe insertion of high density intracortical electrode arrays has been a long-standing practical challenge for neural interface engineering and applications such as brain-computer interfaces (BCIs). Here we describe a surgical procedure, inspired by laser corneal ablation, that can be used in large mammals to thin the pia mater, the innermost meningeal layer encapsulating the brain. This procedure allows for microelectrode arrays to be inserted into the cortex with less force, thus reducing deformation of underlying tissue during placement of the microelectrodes. We demonstrate that controlled pia removal over a small area of cortex allows for insertion of high-density electrode arrays and subsequent acute recordings of spiking neuron activity in sheep cortex. We also show histological and electrophysiological evidence that laser removal of the pia does not acutely affect neuronal viability in the region. This approach suggests a promising new path for clinical BCI with high-density microelectrode arrays.

## Introduction

Intracortical electrodes have been used for decades to study the neural mechanisms of cortical function and disease (Strumwasser, 1958; Hubel, 1959; Hubel and Wiesel, 1959; Bartels *et al*., 2008; Misra *et al*., 2014). More recently, they have also been applied in clinical research as the neural interface component of brain-computer interface systems that restore function after paralysis by connecting the brain to assistive devices or computers (Hochberg *et al*., 2006, 2012; Flesher *et al*., 2016; Lubin, Strebe and Kuo, 2017; Pandarinath *et al*., 2017; Hughes *et al*., 2020). The development of brain-computer interfaces that can record from more neurons (Scholvin *et al*., 2016; Jun *et al*., 2017; Khan *et al*., 2018; Stringer *et al*., 2019; Sahasrabuddhe *et al*., 2020); while minimizing insult to healthy brain tissue (Thelin *et al*., 2011; Kozai, Langhals, *et al*., 2012; Sohal *et al*., 2014; Patel *et al*., 2015; Musk and Neuralink, 2019) is now an active field of research.

While there has been substantial development regarding the electrodes themselves – enabling higher channel counts – there has been little improvement in methods for inserting the electrodes. Inserting microelectrodes into the brains of humans and other large mammals is made difficult by the toughness of their pia mater, a meningeal layer which covers the brain (Wittek *et al*., 2005; Escamilla-Mackert *et al*., 2009; John *et al*., 2017; Weller *et al*., 2018). The force required for an electrode or probe to penetrate the pia mater is often large compared to the resistance of the underlying brain tissue, resulting in deformation (Obaid *et al*., 2019). Consequently, the brain can deform significantly prior to being penetrated by a microelectrode (Obaid *et al*., 2019), and this deformation is empirically associated with tissue damage and loss of neurons over weeks to months (Rousche and Normann, 1992; Kralik *et al*., 2001; Nicolelis *et al*., 2003). This ‘dimpling problem’ is even more pronounced for arrays of electrodes, particularly when the electrodes are closely spaced (Figure 1), (Schwarz *et al*., 2014). The pia mater is nearly impossible to mechanically dissect owing to the presence of intertwined descending blood vessels, which deliver blood to underlying cortical areas (Blinder *et al*., 2013; Shih *et al*., 2013). Therefore, a range of approaches have been adapted for inserting devices *through* the pia mater while minimizing tissue damage.

**Figure 1.**
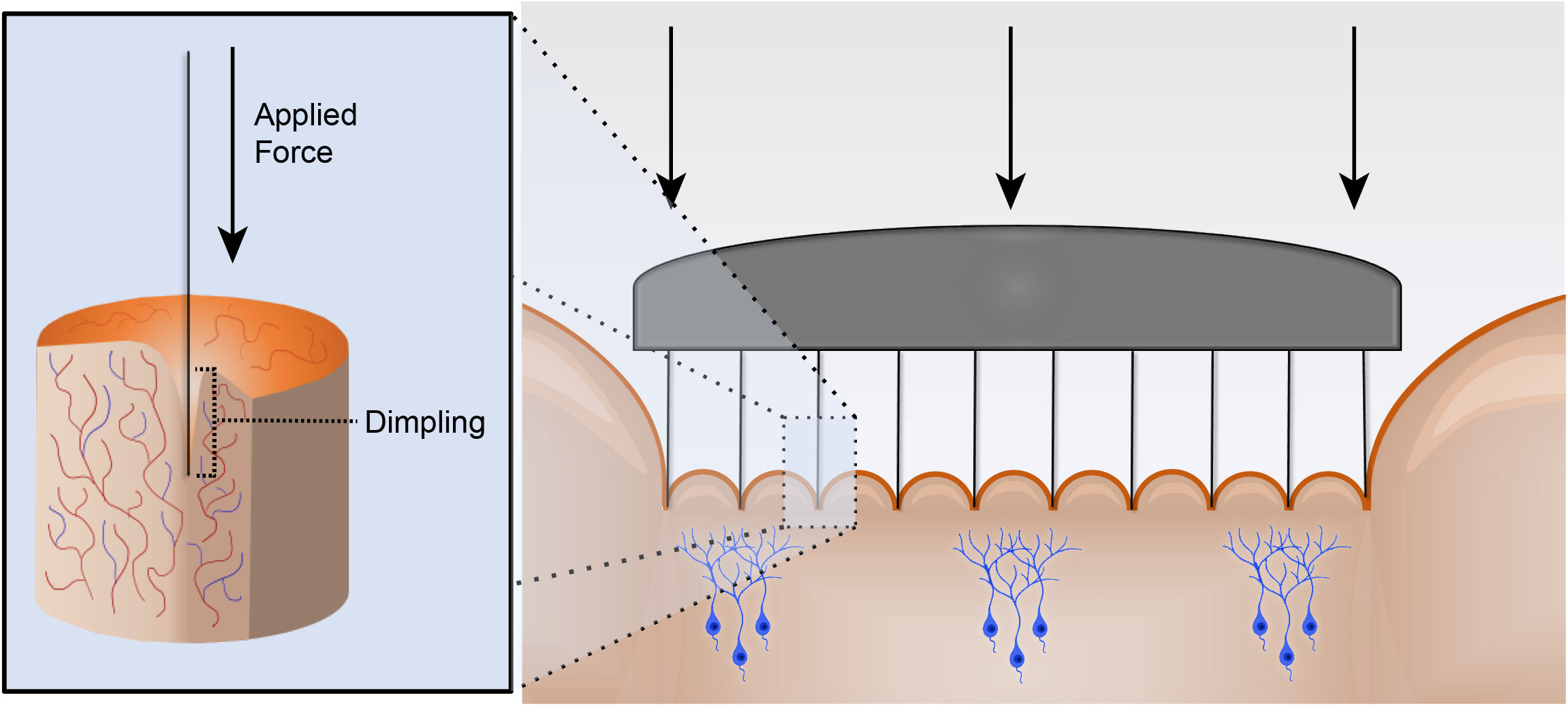
The ‘dimpling problem’. The insertion of microelectrode arrays results in dimpling of cortical tissue because the meninges of the brain resist penetration. An increase in applied force causes greater dimpling of the brain, making high density intracortical arrays difficult to insert. This is compounded by the increased thickness of the meningeal layers in larger mammals and humans.

To overcome the dimpling problem in humans and large animals, the Utah array, which is the current state-of-the-art penetrating microelectrode array deployed for basic and clinical research, is inserted at high velocity (8.3 m/s) (Rousche and Normann, 1992). This high velocity of insertion results in a high strain rate of the underlying viscoelastic brain tissue, resulting in penetration with less cortical deformation than that resulting from slower insertion. However, this implantation strategy can result in bleeding (Csicsvari *et al*., 2003; Barrese *et al*., 2013), and the damage related to insertion has been postulated to be responsible for the variable quality of neuronal recordings in the first few weeks after implantation (Csicsvari *et al*., 2003; Dickey *et al*., 2009; Barrese *et al*., 2013; Vaidya *et al*., 2014).

Another physical approach to electrode insertion is to tension the pia to limit its deformation prior to penetration. This tensioning can be applied by an external mechanical device (Venkatachalam, Fee and Kleinfeld, 1999). A similar effect can be achieved by forcing boundary conditions on the pial membrane by adhesive fixation. Nicolelis and colleagues used an approach where they protect the intended region for insertion of electrodes with the direct application of petroleum jelly to the pia. Cyanoacrylate is then applied around the protected region, with the cured cyanoacrylate holding the pia under tension during subsequent microwire insertion (Kralik *et al*., 2001). However, this technique is limited because the surrounding brain regions cannot be implanted with electrodes due to the cyanoacrylate barrier. Furthermore, if the cyanoacrylate comes in contact with the CSF, this can be detrimental to the targeted brain region and can cause large scale necrosis (Lehman *et al*., 1967; Vinters *et al*., 1985).

Other groups have also implemented chemical removal of the pia using collagenase, either by direct application of collagenase to the pia or by coating the electrode tips with it (Rosenberg *et al*., 1990; Kralik *et al*., 2001; Paralikar, Lawrence and Clement, 2006). The insertion force has shown to be only reduced by 40% (Paralikar, Lawrence and Clement, 2006) and selective removal of the pia confined in the recording region is not always possible, leading to large scale bleeding due to the damage to surrounding vasculature.

An ideal approach to facilitating electrode insertion through the pia mater would be a highly controlled method for pia removal that can be implemented without damaging underlying brain tissue. This would obviate the need for either high speed insertion or physical manipulation of the pia membrane. The pia mater is primarily composed of collagen (Weller *et al*., 2018). In laser corrective eye surgery (e.g. laser-assisted in situ keratomileusis, LASIK), corneal tissue, which is also primarily collagen, is precisely ablated by a pulsed laser (usually 193 nm wavelength excimer laser). This process allows removal of collagenous tissue with submicron precision without causing thermal heating of surrounding areas. The technique has already been successfully used in basic neuroscience research for the removal of protective layers around the brain (Sinha *et al*., 2013). The advantage of an excimer laser is that it can remove tissue without applying significant heat to the tissue (one of the mechanisms is direct disruption of molecular bonds), thereby reducing damage that can be associated with thermal shock mechanisms and scarring of tissue (Vogel and Venugopalan, 2003). Due to the short absorption length of this wavelength and the nanosecond pulse time, this approach does not cause deep tissue damage; non-ablated tissue damage is limited to very thin (<1 µm) layers in the cornea (Vogel and Venugopalan, 2003).

Here we provide evidence that laser ablation of the sheep pia can reduce forces associated with electrode insertion into cortical tissue, both *ex vivo* and *in vivo*, and that single unit recordings can be achieved *in vivo* with a variety of intracortical arrays immediately after laser pia ablation for a period of up to one hour. Histology after pia laser ablation also shows similar neural density to controls for tissues at both the cortical surface and at typical cortical recording depths, demonstrating minimal disruption to neural integrity.

## Methods

### Surgical Approach

This study and all experimental protocols were approved by the Institutional Animal Care and Use Committee (IACUC) at the University of California, Davis, which follows the National Institute of Health (NIH) guidelines for the ethical treatment of animals. White face Dorset Sheep (Ovis aries) that weighed 30 to 35 kg were used for this study, except for the sheep weight vs pia thickness experiments where the weight was explicitly stated. Food was withheld for 24 hours prior to surgery, while water was provided to the sheep *ad libitum*.

Anesthesia was induced using Tiletamine/Zolazepam (Telazol, 4-6 mg/kg, IM). The sheep was intubated, and anesthesia was maintained via 1-5% isoflurane delivered in 60% oxygen and 40% medical grade air. An orogastric tube was placed to minimize or prevent ruminal bloat. Ophthalmic ointment (Paralube, Dechra, UK) was applied to prevent corneal desiccation. Thermal support was provided during the procedure via a circulating warm water blanket (T/Pump, Stryker, USA) or using a Bair hugger (3M, USA).

Once the sterile field was prepared, an incision was made over the skull to expose the bone and underlying fascia. The tissue was reflected, and the periosteum removed over the exposed skull. Next, a surgical microdrill (OmniDrill 35, World Precision Instruments, USA) was used to perform the craniotomy. Bone rongeurs were used to remove excess bone and fully expose the surface of the dura. A craniotomy was made (typically 3 cm × 3 cm) over the motor and auditory cortex in the sheep. Typical stereotaxic coordinates were 5 mm anterior and 25-30 mm lateral from bregma point for the center point of the craniotomy. After the craniotomy, a durotomy was performed using microscissors to expose the pia. The surface of the brain was kept moist with physiological saline throughout the experimental procedure.

Vitals (e.g. SPO2, respiratory rate) were monitored closely during the procedure. To minimize the effects of brain pulsation, end-tidal CO2 was typically maintained between 30-40 mmHg using mechanical ventilation. At the conclusion of the experimental procedures, animals were euthanized with an overdose of sodium pentobarbital (110 mg/kg, IV).

### Laser

A 193 nm, 5 mJ shot energy excimer laser (Excistar, Coherent, Germany) was used to laser-ablate the pia. The set up consisted of a lens tube (Thorlabs, USA) connected to an achromatic lens (Thorlabs, USA) to limit chromatic and spherical aberration of the beam profile. Care was taken to calibrate the power output of the laser with a laser power meter (Thorlabs PM100D, Thorlabs, Newton, NJ) to compensate for the absorption of 193 nm light in air. A galvanic mirror pair was used at the output of the laser to raster the beam across the surface of the brain during the pia ablation process (SCANcube, Scanlabs AG, Germany). A 532 nm targeting laser (Optotronics, Mead, CO) was used to confirm the lased area and to ensure the avoidance of major blood vessels before pia ablation (Figure 2). For all ablations except the force-measurement experiments, a square raster pattern with overlapping 100 parallel lines was used. Ablation areas ranged from 2 mm × 0.6 mm up to 7 mm × 7 mm.

**Figure 2.**
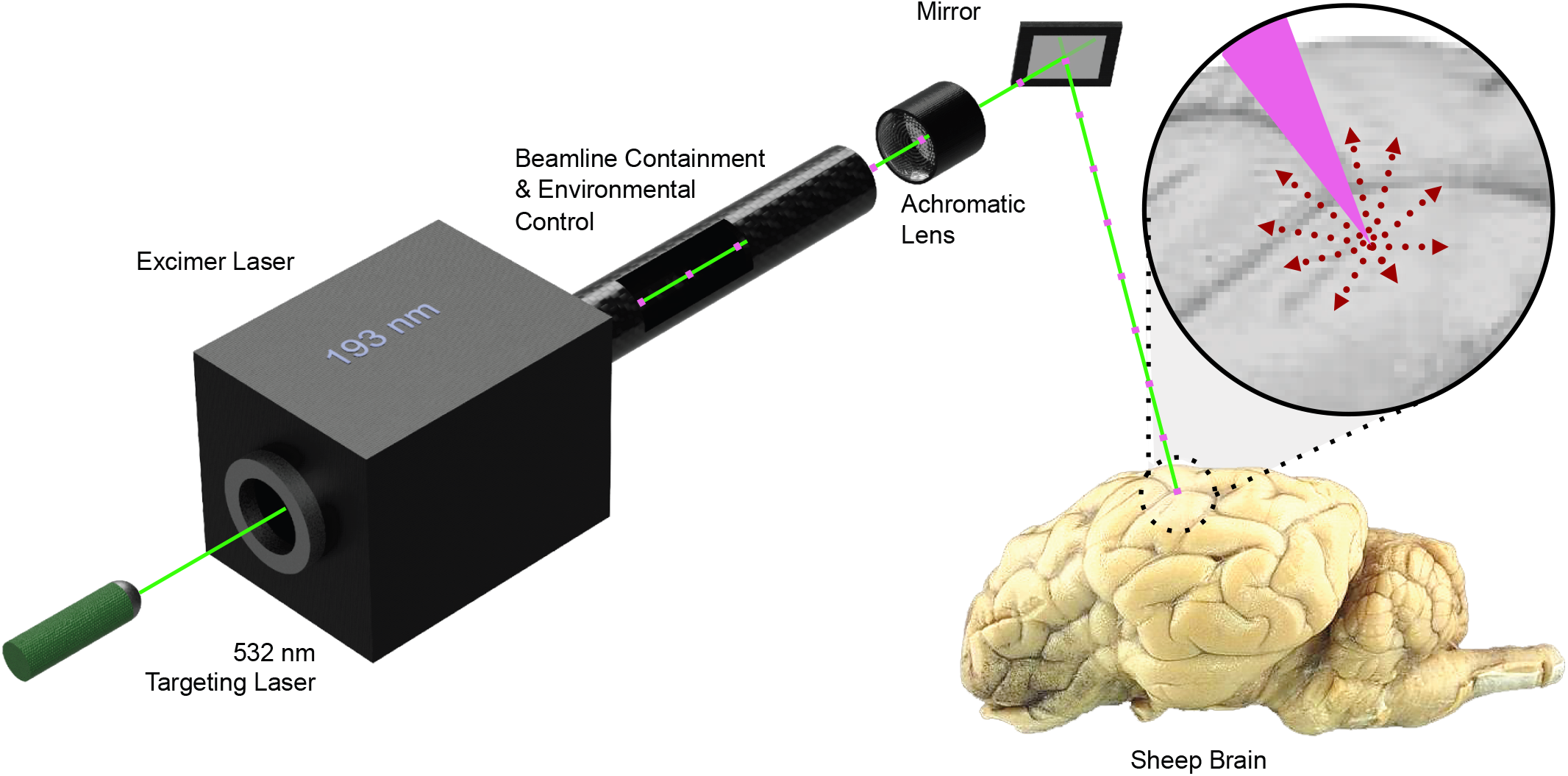
Schematic of the laser pia ablation system. A series of 193 nm excimer laser pulses (purple markers in laser beam path) were used to ablate the pia. Material is removed and ejected from the lased site in a controlled manner (red arrows). Before 193 nm laser ablation, the pre-programmed ablation pattern was evaluated with a 532 nm targeting laser (green laser beam path) to ensure the ablation area was correctly defined and the laser was in focus.

### *Ex vivo* and *In vivo* Force Measurements to Confirm Laser Ablation of the Pia

For the *ex vivo* measurements pieces of brain 2.5 cm × 2.5 cm × 0.6 cm (L × W × D) were excised from the sheep’s cortex immediately (<10 min) after sacrifice of the animal and placed in a petri dish, where they were kept moist. For the *ex vivo* experiments, the lasing was done after excision, while for the *in vivo* experiments lasing was done in situ.

Sites were ablated on the tissue at an energy density of 3.3 mJ/mm^2^ with the number of pulses varied between 0-800. Care was taken that the sites were spaced at least 1 cm apart so that measurements were truly independent. At each site, force measurements were performed and recorded as a function of the number of pulses. In both *ex vivo* and *in vivo* setups, a Futek LSB200 force sensor was mounted (FUTEK Advanced Sensor Technology, Inc, Irvine, California, USA) to a standard stereotaxic manipulator, which had been modified for better cable strain relief. The insertion force was measured using an 80 µm tungsten probe attached to the force sensor. The probe tip was sharpened, and the end subsequently rounded (radius: 30 µm) to measure forces during both the dimpling and penetration phases. The 80 µm electrode diameter and tip shape were both chosen to make measurements more reliable: larger diameter probes result in smaller variation in measured insertion forces, and the hemispherical tip geometry is more reproducible than a sharp tip, in turn making the measurement more repeatable. The probe attached to the force sensor was lowered into the brain at a constant speed of 20 µm/s using an electrically controlled micromanipulator (PI N-381, Physik Instrumente, Karlsruhe, Germany). This movement was stopped once the wire pierced the pia and the measured force started decreasing sharply. Then the wire was retracted, and the setup was moved to the next insertion site.

### Michigan Probe and Utah Array Recordings

Electrophysiology recordings were acquired using the Open Ephys system (Open Ephys, USA). During neural recording, isoflurane was typically reduced to <2% to reduce the effect of anesthesia on neural activity in the brain. The data collected from the system was band-pass filtered between 1 Hz and 7000 Hz at a sampling frequency of 30 kHz. For all recordings, the reference electrode was an 80 μm diameter Teflon poly tetra-fluoro ethylene (PTFE)-coated PtIr wire (AM Systems, USA) One end of this wire was de-insulated for 1-2 mm and placed in the subdural space. The wire was secured in place with moist gel foam (Pfizer, USA).

To record neural activity at various depths, a single shank 32-channel Pt Michigan probe (Neuronexus, USA) was inserted in the cortex in regions with laser pia ablation (n=6) and in control, non-ablated regions (n=4). The 32 channels had 50 μm spacing, distributed over an approximate length of 1.6 mm. The electrode was inserted such that the most superficial site was immediately below the cortical surface. Typical electrode impedance was ∼ 500 kΩ at 1 kHz in physiological saline with an electrode recording site 15 μm in diameter.

For Utah Array recordings (Blackrock Microsystems, USA), a 10 × 10 Pt Electrode Array with 1 mm electrode length and 400 μm electrode spacing was used. The array had a typical impedance of around 200 kΩ at 1 kHz. For recording after laser ablation of the pia, the Utah Array was inserted by manually pushing into the brain with the use of plastic forceps. In one control site, the Utah Array was inserted using a calibrated pneumatic inserter (Blackrock Microsystems, USA). To correctly assess the acute performance of this technique compared to pneumatic insertion, recordings were limited to 1 hour after ablation.

### Histology

Animals were euthanized under general anesthesia with sodium pentobarbital (110 mg/kg). Tissue was collected and placed in 10% neutral buffered formalin and kept at 4°C immediately after euthanasia and dissection to remove relevant brain tissue.

### Cresyl Violet staining for pia thickness measurements

Coronal sections were cut at 50 μm thickness and placed on adhesive glass slides. Care was taken to cut sections from an area where the pia was perpendicular to the coronal plane. The sections were air dried and then baked for 20 minutes at 60°C. Slides were placed in 0.1% cresyl violet (Cresyl violet acetate, Sigma Aldrich, USA) for 5 minutes and staining checked under a microscope, before being dehydrated in absolute alcohol. Sections were cleared with xylene and then mounted for subsequent analysis.

### NeuN staining for Neuronal Density Measurements

For immunohistochemistry (IHC) the formalin-fixed tissue was processed and embedded in paraffin blocks. Coronal sections were cut at 5 μm thickness and placed on adhesive glass slides. The sections were air dried and then baked for 20 minutes at 60°C. Sections were then deparaffinized and hydrated using a series of xylenes and alcohol changes. Antigen retrieval was performed using an Ethylenediaminetetraacetic acid (EDTA) buffer in a steamer followed by rinsing with phosphate buffered saline (PBS).

Primary antibody was applied, and signal amplification was performed using a horseradish peroxidase-labelled detection kit (NeuN antibody #MAB377, MilliporeSigma, USA). Following the application of each solution, it was ensured that the samples were thoroughly rinsed with the use of PBS.

Finally, the Diaminobenzidine (DAB, Sigma, USA) reaction was performed allowing visualization of the primary antibodies and subsequent analysis using optical microscopy. Slides were rinsed with distilled water and counterstained with Gill’s II Hematoxylin (Sigma, USA) to make the pia and superficial layers more visible and allow for pia thickness measurements. Slides were dehydrated and cover slipped for permanent mounting.

### Image Analysis

Slides were scanned with an Olympus PlanApo 20x/.40 NA objective and a global shutter, CMOS sensor camera paired with a .63x adapter. Once the slides were scanned, Aperio ImageScope (v12.4.0.5043) was used to image the region of interest (ROI) at a 10X magnification. For cresyl violet staining, the pia thickness was measured using a calibrated scale bar through collected images.

For NeuN staining, images were collected across two regions of interest in the cortex, the surface (100-400 µm deep, ROI: 300 µm × 1 mm) and recording (900-1000 µm deep, ROI: 100 µm × 1 mm) depths across samples. This allowed the assessment of the number of neurons in regions close to the lased pia and at a depth where the recording tips are likely to reside in electrode implants most commonly used in basic neuroscience and clinical research. Control samples (no laser ablated pia) were taken from the contralateral hemisphere from the same cortical region. The images were saved and NeuN stained cells were counted using the ImageJ “Multi-point” tool. Analysis was performed and compared between lased and non-lased (control) tissue samples. At least 3 sections for each site were taken, and cell counts were normalized with the ROI area to provide a metric of neurons/mm^2^.

### Offline Recording Analysis

Neural signals recorded by Open Ephys were imported into MATLAB for analysis (MATLAB 2018b, Mathworks, USA). Channels were appropriately filtered (using ‘filtfilt’ function to avoid phase distortion) from 300-6000 Hz to obtain the spiking frequency band of interest.

Spike sorting was performed using Wave_Clus (Quiroga, Nadasdy and Ben-Shaul, 2004). Typical thresholds for neural data were set at least 3.5 times the noise threshold, calculated through Wave_Clus. In short, the noise threshold (σ) was calculated by taking the absolute mean of the bandpass signal and dividing it by 0.6745, and the spike crossing threshold was set at a minimum of 3.5σ (Quiroga, Nadasdy and Ben-Shaul, 2004). The output of this batch processing was then used to confirm the presence or absence of spike waveforms. Single unit activity was confirmed using three metrics: (1) all neural waveforms had a peak width less than 1 ms. (2) The interspike interval histogram (Quiroga, Nadasdy and Ben-Shaul, 2004; Rossant *et al*., 2016) showed a clear indication of a refractory period (i.e. no waveforms in the 0-3 ms bins on Wav_Clus output). (3) Clusters were clearly separated as confirmed through the Wave_Clus user interface.

### Statistical Analysis

Statistical tests were performed in MATLAB (MATLAB 2018a, Mathworks, USA) and Python. Unless otherwise mentioned, the 95% confidence interval was estimated as 2 times the standard error of the mean. For all statistical tests performed, results were deemed significant if *P<0*.*05*.

For the pia thickness measurement, 10 sheep were evaluated. For four weight conditions n=2 sheep were evaluated each and for the two heaviest weight conditions n=1 sheep were evaluated each. For each sheep, 3 sections were taken and analyzed. When applicable, the data over sections and animals was pooled for calculation of the standard error of the mean.

For force measurement comparisons between *ex vivo* and *in vivo* tissue, Mann-Whitney U tests were performed and compared for 0, 50 and 400 pulses at 3.3 mJ/mm^2,^ with the insertion force for each modality having been measured 3 times.

To compare the total number of neurons for Michigan probe recordings between control and pia ablated brains, a Mann-Whitney U test was performed over the average number of units per recording site. To compare the effect of lased site area (2 mm × 0.6 mm, 3 mm × 3 mm and 7 mm × 7 mm) on the number of neurons obtained, the number of units was pooled over repeat experiments and a Kruskal-Wallis test was performed. To compare the spatial variation in unit count across the Utah Array, central (n=36 per array) and edge electrodes (n=60 per array) from the Utah Array were analyzed. Data were pooled from both recordings, yielding 72 samples for the central area and 120 samples for the edge areas, totaling 192 samples.

For the NeuN staining analysis, comparisons were performed using percentile bootstrapping (Sohal *et al*., 2014; Falcone *et al*., 2020) due to unequal sample sizes between control sites (n=5 sites with 15 sections) and lased sites (n=13 sites with 39 sections). Total number of neurons/mm^2^ from the full dataset were drawn randomly (with replacement). This procedure was repeated 10,000 times to estimate the expected difference in each measure. The null hypothesis for this measurement was that there is no difference in number of neurons/mm^2^ between control and lased site samples. The null hypothesis was rejected if the observed difference fell outside the 2.5– 97.5% percentile range of the bootstrapped distribution.

## Results

### Validation of the sheep model for human translation

Structurally, the sheep brain is very similar to the human brain in terms of brain folding (i.e. sulci and gyri formation) (Figure 3a). To further ensure that the sheep cortex was a valid model to test the laser system and insertion dynamics of electrodes we compared the pia mater in sheep and humans (Figure 3b) and found that the average sheep pia thickness was 76.8±3.8 µm (±SEM, n=30, Figure 3c). This is in good agreement with human pia thicknesses, which varies between 50 and 100 µm (Wittek *et al*., 2005; Escamilla-Mackert *et al*., 2009; John *et al*., 2017; Weller *et al*., 2018). Therefore, the sheep cortex is a suitable model for testing intracortical electrode insertions and understanding pia laser ablation dynamics.

**Figure 3.**
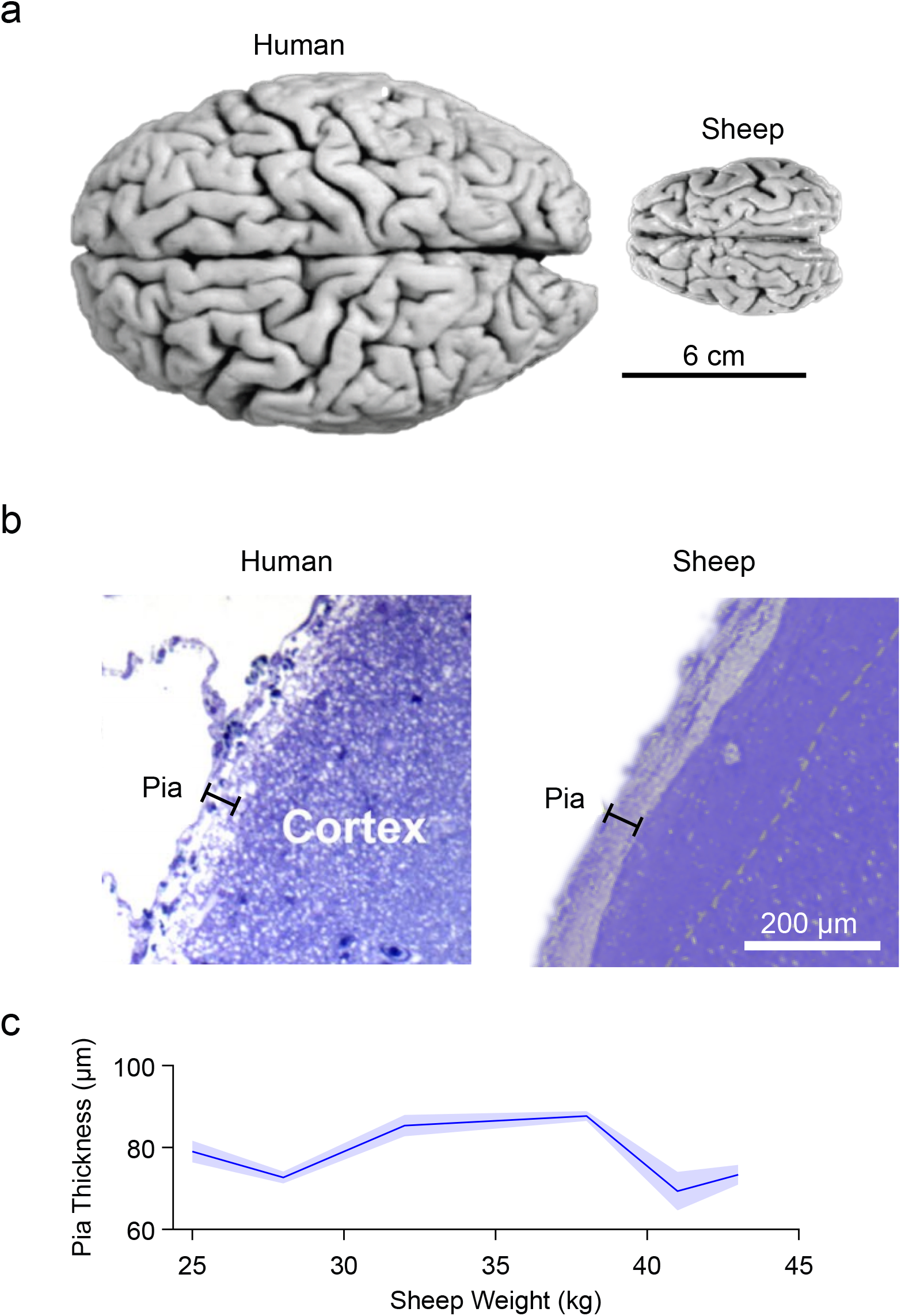
A comparison of sheep and human brain. (a) The sheep (ovine aris) brain has sulci and gyri similar to those of the human brain. (b) A histological comparison of the pia and cortex between human (Weller et al., 2018) (left) and sheep (right, cresyl violet, n=10 sheep) demonstrates the similarity in pia thickness. (c) In sheep weighing 25-45 kg, the thickness of the pia is consistently 70-80 µm (mean ± SEM shown), and similar to the thickness of the human pia (50-100 µm).

### Laser ablation of the pia mater reduces insertion force in a dose-dependent manner

We used a custom probe setup with an attached 80 µm tungsten wire to measure insertion force while applying an increasing number of laser pulses (0 to 800) at an energy density of 3.3 mJ/mm^2^ (Figure 4a). We found that after an initial period of negligible force reduction (0 and 25 shots: 31.5±5.2 mN, n=6, ±SEM) - which we interpreted as removal of water and other contaminants from the surface - there was a roughly linear relationship between applied laser pulses and insertion force. These two factors are likely connected through remaining pia thickness as a mediator variable. After 400 shots, the force had dropped to 0.28±0.06 mN (n=3, ±SEM). Doubling the number of pulses from 400 to 800 did not yield further reduction in force, consistent with the assumption that at that point the pia had been entirely removed from the intended location.

**Figure 4.**
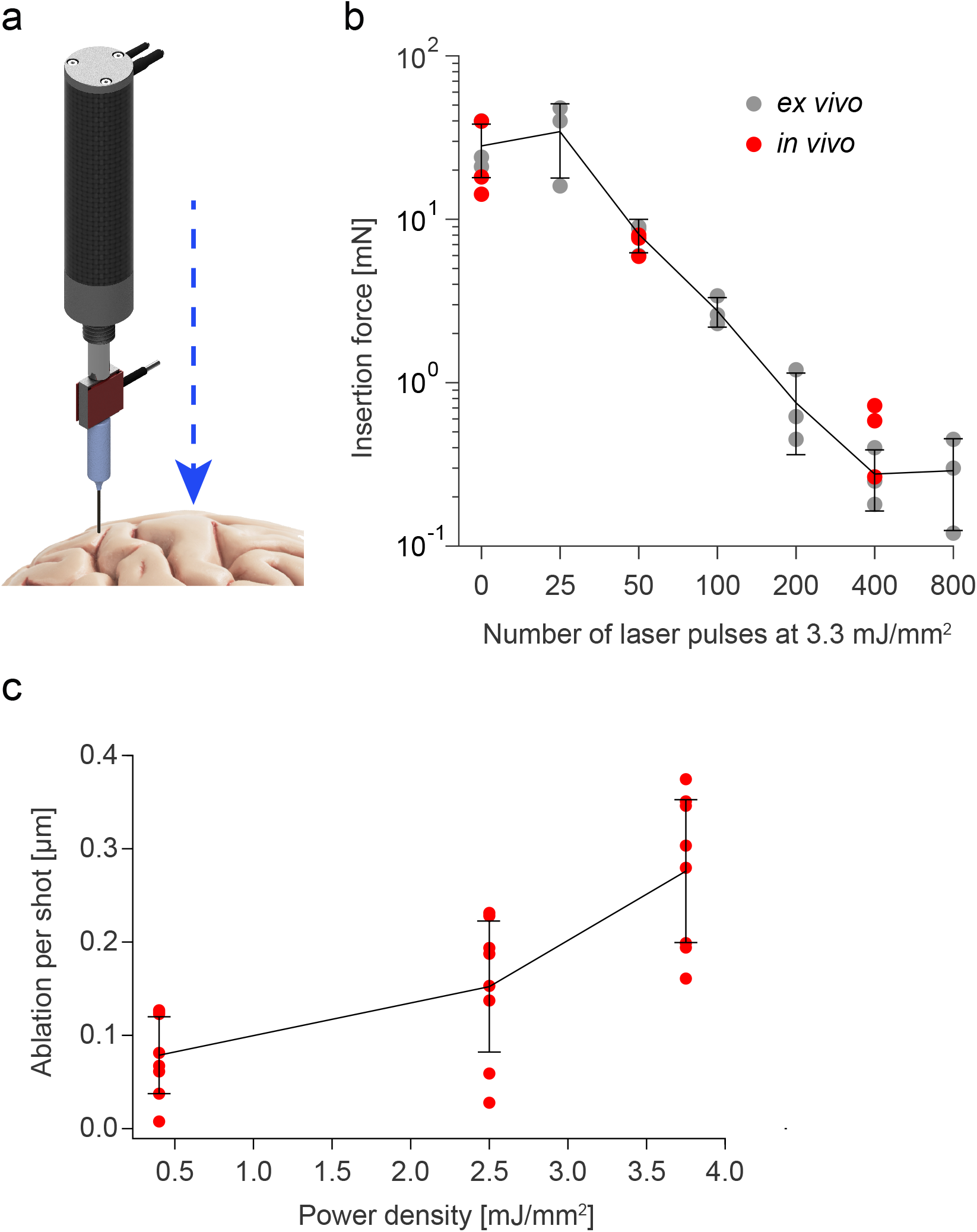
Confirmation of pia laser ablation *ex vivo* and *in vivo*. (a) A schematic of the probe set up used to acquire force measurements. Shown are the actuator (black), the force sensor (red) and the probe itself approaching the brain. (b) Insertion force as a function of number of pulses (0 and 25 to 800 pulses plotted on a log scale) per site at an energy density of 3.3 mJ/mm2 per shot. Mean and standard deviation shown as black line. The decrease in insertion force follows the same trendline with similar ablation parameters for both *ex vivo* and *in vivo* tissue. (c) *In vivo* Pia laser ablation as a function of power density. Sites from the brain were laser ablated at different power densities and pia thickness was determined through coronal sections (n=8 per power density, mean and standard deviation shown as black lines). Increasing the power density increases the amount of pia ablated in a controlled manner.

There was good agreement between force measurements as a function of laser pulses between the *ex vivo* and *in vivo* experiments (Figure 4b). No significant differences were found for force measurements from 0, 50 and 400 laser pulses between *ex vivo* and *in vivo* measurements (P>0.20, P>0.40, P>0.40 respectively, n=3, Mann-Whitney U test)). We then assessed the minimal power to reliably ablate the pial tissue and found that power densities as low as 0.4 mJ/mm^2^ can ablate the pial tissue (Figure 4c). A reduction in shot energy can be desirable to reduce percussive stress to the tissue (Vogel and Venugopalan, 2003).

In summary, pia laser ablation is an effective tool for reducing insertion force for cortical electrodes and can be accurately controlled both by the number of lasing pulses and energy deposited per pulse.

### Pia Laser Ablation Does Not Inhibit Neural Recordings

#### Michigan Probe Recordings

To assess neural viability after lasing, a multi-depth, single-shank Michigan probe was inserted in the middle of the target area immediately after laser ablation of pia. The probe was inserted so the top-most recording site was just in the brain to remain superficial in the cortex. Across recordings in laser-ablated regions (n=6) as well as in control locations (n=4), we isolated single units along the shank (Figure 5a) and found that the average number of neurons per site in the lased condition was 0.93±0.13 (±SEM, n=6) and 0.81±0.18 (±SEM, n=4) in the control condition. This difference was not significant (P>0.13, Mann-Whitney U test). Furthermore, even among individual recording sites the difference between the lased and control condition was not significant (minimum significance P>0.10 at site 20, Mann-Whitney U test) (Figure 5b).

**Figure 5.**
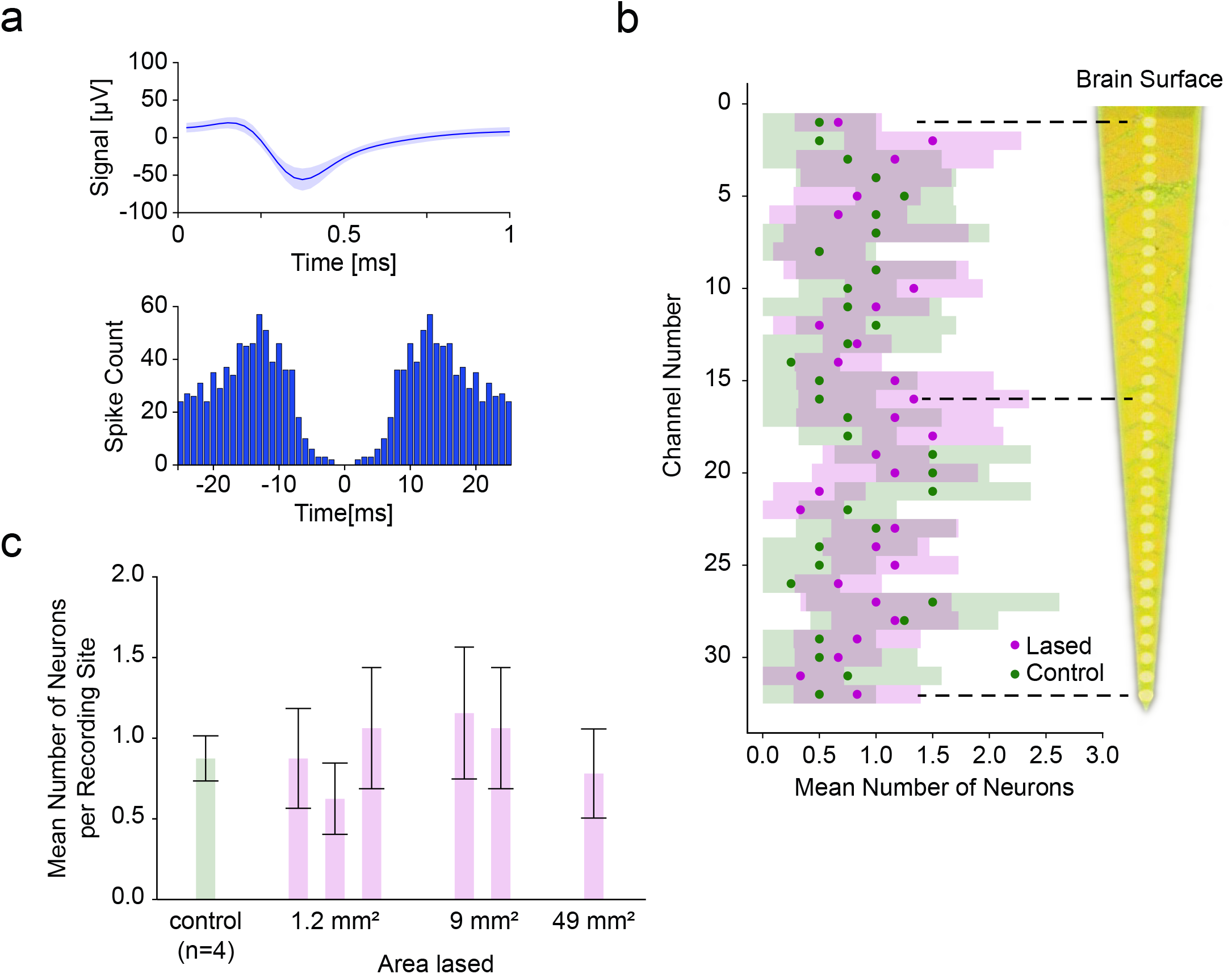
Comparison of single-shank Michigan probe depth recordings between control and laser ablated sites. (a) The mean spike waveform (±SEM) and autocorrelogram for one of the recorded single units. (b) Average number of neurons (±95% CI) recorded at each depth was not significantly different between lased tissue (n=6) and control tissue (n=4) (most significant at site twenty: P>0.10, Mann-Whitney U test). The distance between recording sites was 50 µm.(c) Average number of neurons (±95% CI among recording sites) as a function of increasing the lased site area. No significant differences in neural count were found (P>0.10, Kruskal-Wallis test), showing the ability to enlarge the lased site area without detrimental effect to the number of isolated neurons.

Next, we investigated whether the size of the ablation would have an influence on the recorded unit count. We therefore sorted the data from Figure 5b by lasing area and found that for areas 2 mm × 0.6 mm (n=3), 3 mm × 3 mm (n=2) and 7 mm × 7 mm (n=1) the average number of units per recording site was 0.85±0.08, 1.11±0.11 and 0.78±0.12, respectively, (Figure 5c, ±SEM, n=96, n=64, n=32 units; n=3, n=2, n=1 recordings respectively) which was not a significant difference (P>0.10, Kruskal-Wallis test). This shows the viability of pia laser ablation to accommodate the size footprints of most commercial arrays (e.g. the Utah Array’s active electrode area of 4 mm × 4 mm) without a detrimental effect on acute neural function.

#### Utah Array Recordings

To demonstrate the compatibility of our lasing approach with multielectrode arrays insertion and recording, we chose a commercially available intracortical electrode array, the Utah Array. Following laser ablation of pia over a 6 mm × 6 mm area, the array was inserted into cortex using forceps, and neural activity was recorded (Figure 6a).

**Figure 6.**
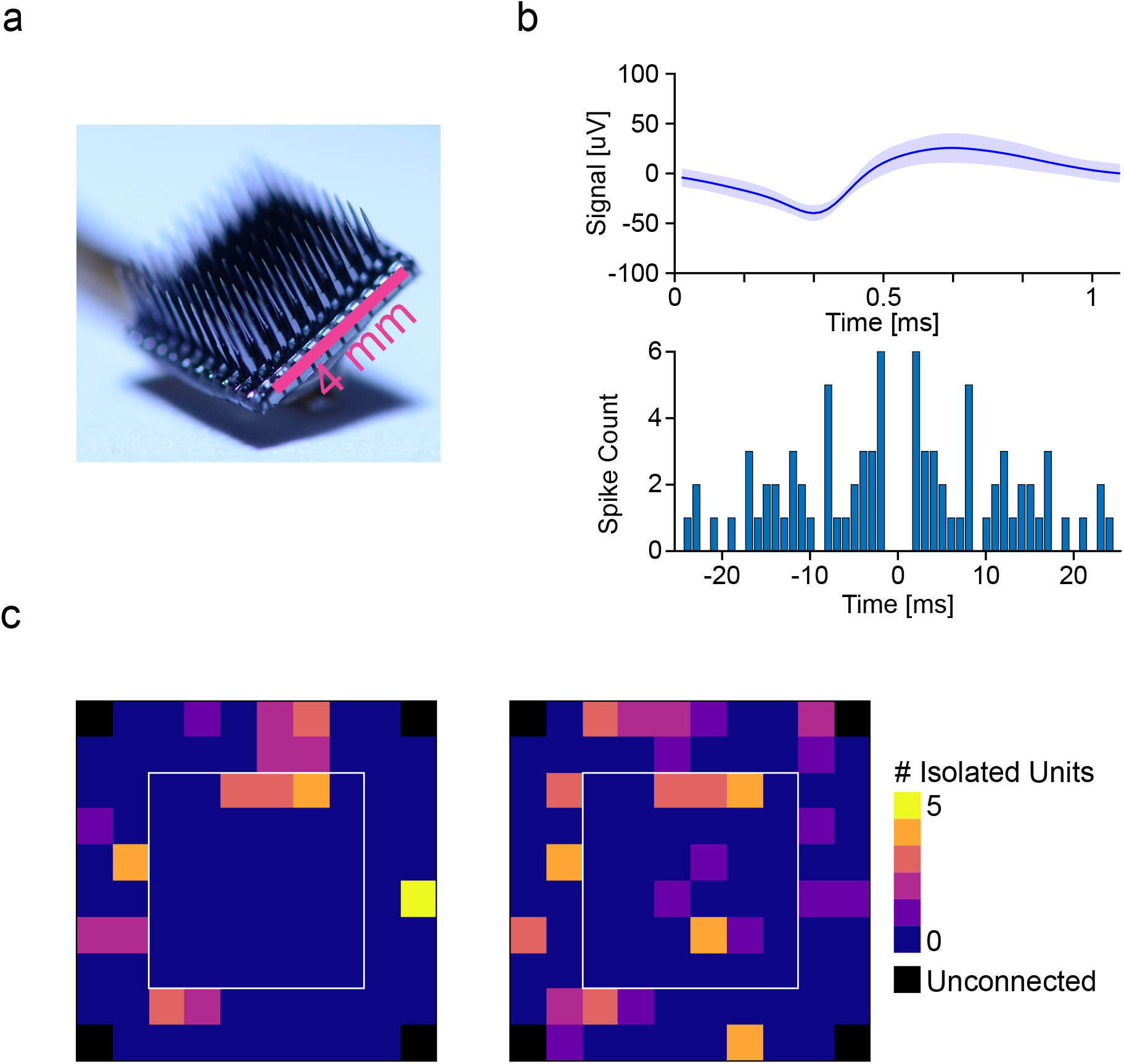
Comparison of Utah Array (a) recording from laser ablated manual insertion. (b) Representative overlain isolated unit from Utah Array recording with a clear formation of an autocorrelogram, confirming unit isolation. (c) 2D plots (top down view) of the Utah array electrode grid for two recording sessions, showing number of isolated units. The number of isolated units in the center area (white frame) was not significantly different from the outside area (see results), suggesting (for a lased area of 6 mm × 6 mm) the absence of edge effects in the lasing viability evaluation.

Results from two inserted Utah Arrays following pia ablation show the ability to record units immediately after laser ablation in the sheep cortex (Figure 6b). In these two experiments, we found that directly after insertion of the arrays we were able to record from 39 and 53 isolated single units, respectively. For comparison we also inserted a Utah Array into sheep without lasing (using the pneumatic inserter) which resulted in 21 isolatable single units (n =1).

The spatial trends of unit count across the Utah Array were also investigated to rule out specific areas of damage to the cortex with laser ablation (Figure 6c). The outer 60 electrodes had a mean single unit yield of 0.54±0.12 (±SEM, in aggregate over both recordings, n=120), compared to 0.38±0.10 for the central 36 electrodes (±SEM, in aggregate over both recordings, n=72) and that there was no significant difference between those two regions (P>0.23, tested in aggregate over both recordings, Mann-Whitney U test).

### Histology Shows No Detrimental Neural Loss After Pia Ablation

To compare the viability of neurons for control and laser ablated sites, NeuN staining was performed through IHC protocols with the number of neurons/mm^2^ calculated from coronal sections. For analysis, the electrode sites along the shank were divided into two regions of interest: the proximal electrode sites which recorded superficially at a depths of 100 µm to 400 µm below the cortical surface (orange region in Figure 7a-b) and the distal sites, which recorded at depths of 900 µm to 1 mm below the cortical surface (blue region in Figure 7a-b). No significant differences in neural density were observed (surface layers: 111±49 (control), 121±41 (lased), recording depth: 340±139 (control), 329±135 (lased), all numbers mean ± 95% CI, Figure 7c) between control (n=5) and lased sites (n=13) for both regions of interest (percentile bootstrap, surface layers P>0.09 and recording depth P>0.29). Thus, laser ablation of the pia does not detrimentally affect neural density in an acute setting at the recording depth regions and with no superficial damage to the sheep cortex.

**Figure 7.**
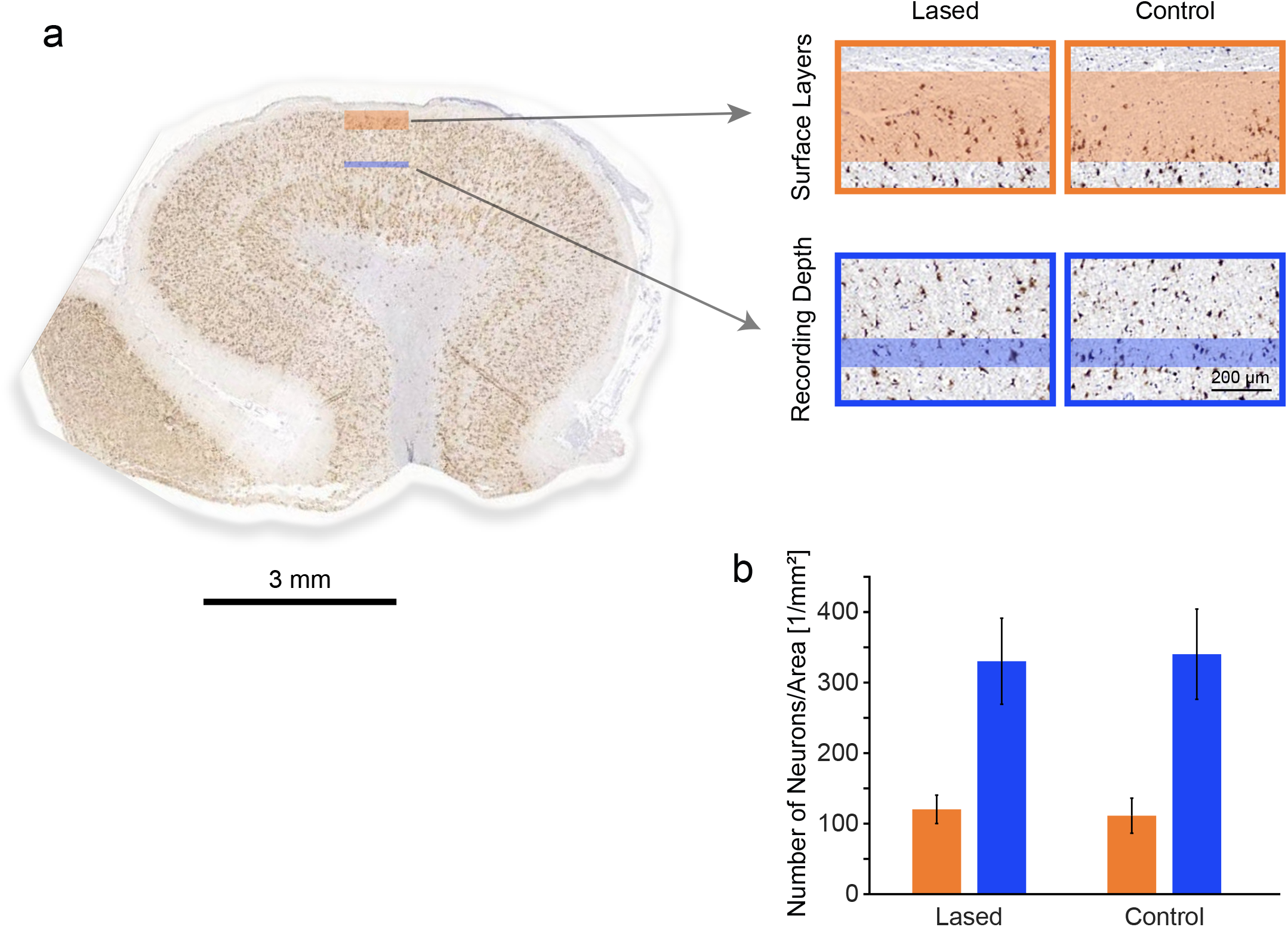
Histological comparison of pia tissue between control sites (n=5 sites) and lased sites (n=13 sites) for superficial and recording depth profiles. (a) Coronal section showing measurements locations for neuron density (neurons/mm^2^) through NeuN staining were taken for both superficial (orange box) and depth (blue box) recording profiles. (b) Representative coronal sections for laser ablated sites and control sites for superficial (orange box) and recording depth (blue box) profiles. (b) Mean (±95% CI) number of neurons/mm^2^ for laser ablated and control sites for both depth profiles. The number of neurons between laser and control sites are comparable for both depth profiles, demonstrating a lack of acute damage to neurons through laser ablation of the pia (percentile bootstrap, P>0.09 and P>0.29 for surface layers and recording depth, respectively).

## Discussion

Here we provide compelling evidence that controlled laser ablation of the pia can facilitate insertion of commercial microelectrode arrays into the cortex of a large mammal (sheep) without causing acute harm to tissue. All arrays used in this study had high unit count with laser ablated pia sites, and we observed typical neural density at both surface and recording depth profiles compared to control brains.

### Sheep model

In this study we used sheep to study changes in insertion dynamics after using laser ablation of the pia. The use of large animal models in translational BCI research and development is essential, because the biomechanics of smaller brains and their associated meninges will differ substantially from humans. Historically, microelectrode studies for BCI applications have been primarily performed in non-human primates (NHPs). However, the behavioral training required for NHP studies is time consuming (Taylor, 2002; Fraser and Schwartz, 2012), and NHPs are expensive and heavily regulated in the United States and elsewhere. Thus, studies are practically limited to a small number of animals over the course of a year. Due to these practical restrictions, it was highly desirable to perform early device testing in an alternative large mammal.

The sheep brain has a similar folding to the human brain and the pia has a similar thickness to that of humans, thus making the sheep an appropriate model system for understanding the dynamics of intracortical array insertion. Moreover, sheep have similar cortical architecture to humans, and have been used in a variety of translational studies (John *et al*., 2017; Opie *et al*. 2018), making sheep also suitable as a translational model for electrophysiological medical devices.

### Force Measurements and Dimpling

In our study, force measurements were used to validate the thinning of the pia. For non-lased samples, force measurements followed the same pattern as those previously reported, where dimpling of the brain was reported to be 1-2 mm in a mouse model using an 80 μm tungsten wire probe, inserted at 20 µm/s (Obaid *et al*., 2019). Interestingly, the dimpling of the sheep brain could exceed 3 mm under the same conditions. The dimpling depth is a monotonic function of the force needed to insert so it is possible to use dimpling depth as a rough proxy for insertion force when it is not feasible to measure insertion force. Techniques for reducing this force and subsequent dimpling include thinning the pia and working towards thinner probes (Thelin *et al*., 2011; Kozai, Langhals, *et al*., 2012; Guitchounts *et al*., 2013; Patel *et al*., 2015; Obaid *et al*., 2019) as we approach translational brain models and human pia thicknesses.

### Michigan Probe Depth Recordings

Michigan probes were used in this study to obtain an electrophysiological depth profile of the neural layers of interest following laser ablation of the pia. If there was damage to specific underlying cortical regions, the expectation would be the lack of neurons at certain depths along the shank of the array. As we obtained similar unit count across depths for laser ablation in comparison to control, it is likely that pia thinning had little effect on compromising neural function. Overall, yields of 0.93 units per site were found for our Michigan probe recordings, similar to acute recordings using Michigan probe-style electrode arrays from other animal models (Ludwig *et al*., 2009; Du *et al*., 2011; Jun *et al*., 2017; Csicsvari *et al*., 2003).

### Utah Array Recordings

Typically, Utah Arrays are inserted at high velocity using a pneumatic inserter to overcome the dimpling issue of the meninges (Rousche and Normann, 1992). In this study, the Utah Array can be inserted into the cortex with a simple forceps approach following laser thinning of the pia. We found that the unit count in our initial Utah Array recordings with pia ablation-assisted insertion exceeds the unit count we found directly after pneumatic insertion (39 and 53 vs. 21). This highlights the utility of the lasing paradigm as a tool for acute Utah Array recordings.

### Histology

In this study we used NeuN staining to measure neural density for control and pia laser ablated sites. In keeping with the electrophysiological analysis, we analyzed both superficial and deep regions of cortex to assess if areas closer to the ablation zone were differentially affected. NeuN was used as it was a directly available measure of neural integrity over the course of the experiment. Typical experiments were 8 hours, with all tissue samples collected post-euthanasia. Therefore, initial long-lasting effects of pia ablation would have been captured over this time course during the acute experiment. However, even for these long time periods after lasing, the neural density in pia -ablated sites was still comparable to control brain samples.

Although we provide compelling electrophysiology and histology showing the viability of pia laser ablation to aid insertion of commercial arrays, long-term studies are still needed to evaluate the method. In this acute study we were limited to NeuN staining to evaluate the neural density for both control and pia laser ablated sites. It is well known that typical markers of the glial response (e.g. microglia and astrocytes) are either not activated or seen to migrate to the site of electrode implantation during the time duration of an acute experiment (Kozai, Vazquez, *et al*., 2012; Wellman *et al*., 2019). However, in long-term implantation, other relevant markers for gliosis (e.g. ED-1-microglia, GFAP-Astrocytes, IgG-blood brain barrier integrity) can be used to determine the integrity of the tissue (Polikov, Tresco and Reichert, 2005; Nolta *et al*., 2015; Sohal *et al*., 2016; Falcone *et al*., 2019). This combined with chronic electrophysiology and impedance measurements will be indicative of the electrode performance with laser ablation of the pia used as a method for insertion.

### Future Directions

Here we describe the acute electrophysiological and histological effects of pia removal. To fully evaluate the promise of laser ablation of the pia as an insertion method, electrophysiological validation should be extended to chronic recording and histology performed weeks or months after implantation. Animals should be chronically implanted with commercial arrays and compared to appropriate controls for pia laser ablation and control brains with non-ablated pia using conventional insertion methods.

Due to the high shot rate of the laser used in this study all ablations could be performed in seconds. The time consumed by the lasing was dominated by the alignment of the lasing setup with the desired target region. This time was kept low by having a dedicated laser technician present during all experiments. For a broader applicability it would therefore be desirable to have a more integrated system available that can reduce the setup time and automate the alignment task.

Although the laser was able to safely ablate areas as large as 7 mm × 7 mm, it was evident that attempting a larger-sized ablation would risk damaging the walls of prominent vessels, causing them to rupture to the detriment of surrounding neural tissue. A future system would be greatly enhanced by combining patterned laser ablation with blood vessel imaging and detection. By registering the vessel map with the scanning ablation beam, larger areas could be exposed without risking major bleeding. This kind of vessel avoidance is a common method that is already employed in various studies including those for intracortical electrode insertion (Takahashi *et al*., 2000; Markwardt *et al*., 2017; Ramakonar *et al*., 2018; Musk and Neuralink, 2019).

## Acknowledgements

This work was funded by the Defense Advanced Research Projects Agency’s Neural Engineering System Design program (N66001-17-C-4005).

We would like to thank Leigh Hochberg, MD, PhD, Sydney S. Cash, MD, PhD, Vikash Gilja, PhD and Angeles Salles, PhD for comments on the manuscript, Mina Hanna, PhD for fruitful discussions, Alison Stuckey, Yeena Ng, Sashank Shivakumar and Devin Fell for outstanding technical and logistical support, and Marike Zwienenberg, MD, Linda Talken, and Amy Lesneski (UC Davis) for help in developing sheep surgical protocols.

## Competing interests

All authors are current or former compensated employees of Paradromics, Inc., a brain-computer interface company.

